# Phase transitions and symmetry breaking of cooperation on lattices

**DOI:** 10.1101/2025.03.24.644969

**Authors:** Christoph Hauert, György Szabó

## Abstract

The donation game is an instance of a social dilemma with a single parameter given by the cost-to-benefit ratio of cooperation, *r*. In spatial settings limited local interactions and clustering are capable of supporting cooperation by reducing exploitation from defectors. Traditionally the interaction and competition neighbourhoods are identical. Here we discuss intriguing differences in the dynamics that arise when separating the neighbourhoods. On the square lattice disjoint interaction and competition neighbourhoods are easily realized by considering nearest neighbour interactions and second nearest neighbour competition. Incidentally, the number of first and second neighbours is the same. More importantly, this separates the population into two competing sub-populations, with interactions solely between sub-populations but competition within sub-populations. For negative cost-to-benefit ratios, *r*, the donation game turns into a harmony game and defection becomes an act of spite. In the traditional setup the extinction of cooperators under harsh conditions, large *r*, and that of spiteful defectors, *r <* 0, exhibits critical phase transitions with characteristics of directed percolation. In contrast, with two sub-populations spiteful behaviour cannot persist, while the extinction of cooperators exhibits the same characteristics. Most intriguingly, however, for smaller *r* spontaneous symmetry breaking in the levels of cooperation between the two sub-populations is observed. The symmetry breaking resembles the sub-lattice ordering occurring in the anti-ferromagnetic Ising model. Within the twofold degenerated phases, decreasing the cost-to-benefit ratio induces extremely large fluctuations (bursts) in the frequencies of cooperation. These bursts eventually drive the system into one of the absorbing states: occasionally homogeneous defection in both sub-lattices but usually only in one and homogeneous cooperation in the other, achieving perfect asymmetry.

**PACS numbers:** 89.65.-s, 89.75.Fb, 87.23.Kg

## I. INTRODUCTION

Concepts and methods from statistical mechanics have proven instrumental for the exploration and quantification of a wide range of dynamical phenomena in evolutionary game theory [1–9] and the evolution of cooperation in spatial settings, in particular [10–12]. In spatial models players are arranged on a lattice or represented by nodes in a network. Through interactions with their neighbours each individual obtains a payoff. The payoffs indicate the success of this individual in terms of higher chances for producing clonal offspring or serving as a model for imitation by others. In either case, successful strategies have higher chances to increase in abundance than less successful ones. This mimics genetic or cultural evolutionary processes.

In evolutionary games a (probabilistic) update rule determines how players get replaced by the offspring of more successful individuals or, equivalently, how they modify their own strategy through comparisons of their own performance to that of others [13, 14]. In spatial settings the stationary state and strategy distribution of such evolutionary processes is determined by the payoffs of interactions, the neighbourhoods of players for interactions and for competition (imitation or reproduction), as well as the microscopic updating of individual strategies. The following sections explain in detail how to assign payoffs to individuals, mimic spatial structure through interaction and competition neighbourhoods as well as how to characterize the stationary states.

### A. Interactions

The simplest class of interactions is given by 2 × 2 games, where two players meet and have two strategies at their disposal. The most prominent example is the prisoner’s dilemma, which captures the conflict of interest between the performance of either player and the pair of them. This conflict of interest represents the hallmark of social dilemmas [15, 16]. A particularly simple instance of the prisoner’s dilemma is given by the donation game [17]: a player can *cooperate, C*, and provide a benefit, *b >* 0, to their partner at a cost, *c < b*, or *defect, D*, and do nothing. If both players cooperate they each end up with *b* − *c >* 0, which is preferable over a payoff of zero for mutual defection. However, each player is always tempted to defect because defection pays more than cooperation regardless of the partner’s behaviour – to the detriment of all. As a consequence defection is a dominant strategy and mutual defection the sole Nash equilibrium with zero payoff. At the same time both players prefer mutual cooperation *b* − *c >* 0.

The donation game is conveniently summarized in the form of a payoff matrix (for the row player):

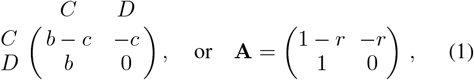

with the cost-to-benefit ratio *r* = *c/b*. Without loss of generality the payoffs can be rescaled by dividing by *b*. This results in an intuitive, single parameter description of the interaction. For *r >* 0, the social dilemma applies and defectors are better off regardless of the opponents decision. This is a consequence of the ranking of the four payoffs, which determines characteristics of the interaction. In contrast, for *r <* 0 the ranking changes and the social dilemma essentially disappears. The only remnant of the social dilemma is that defection harms the partner more than the defector itself, who is thus better off. This is the definition of spiteful behaviour [18, 19]. At the same time it is now better to cooperate regardless of the opponents decision. The resulting interaction is often termed a harmony game because now cooperation is the dominant strategy. Mutual cooperation is the preferred outcome and the sole Nash equilibrium. Finally, for *r* = 0 the payoff to an individual is independent of its own strategy and hence exploitation (as well as spiteful behaviour) is impossible. In a spatial setting this recovers the voter model [20].

An alternative and widely used single parameter description is given by the *weak* prisoner’s dilemma [10] with the payoff matrix:

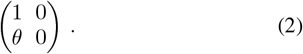

It is called weak because cooperation entails no cost and hence cooperators and defectors perform equally well against defectors. As a consequence mutual defection turns into a *weak* Nash equilibrium. With the benefits for mutual cooperation normalized to one, the only remaining payoff is the temptation to defect, *θ >* 1. Thus, the payoffs defy the traditional classification and characterization of interaction types based on their ranking. Instead, they refer to the border between prisoner’s dilemma [21] and hawk-dove [1] or snowdrift games [22]. In spatial structures the expectation for the two games differ: Limited local interactions invariably benefit cooperators in the prisoner’s dilemma but may inhibit cooperation in the snowdrift game [23, 24]. For 0 *< θ <* 1 the weak prisoner’s dilemma again turns into a (weak) harmony game with the potential for spiteful behaviour.

### B. Spatial Games

Based on the characteristics of the donation game it is clear that cooperation is doomed in the absence of supporting mechanisms. A well-studied situation that is capable of supporting cooperation is to include spatial dimensions. Through limited local interactions and cluster formation cooperators are able to reduce exploitation [10] and take over the population [25] or co-exist with defectors in a dynamical equilibrium [26] – at least under sufficiently benign conditions as reflected by small cost-to-benefit ratios *r* in Eq. 1.

In the simplest case players are arranged on a square lattice with linear extension *L* and *N* = *L* × *L* sites. Boundaries are periodic to eliminate boundary effects and to reduce finite size effects. Every player interacts with its *k* = 4 nearest neighbours (von Neumann neighbourhood). The strategies to cooperate, *C*, or defect, *D*, for a player located at site *i* can be encoded as two-dimensional unit vectors

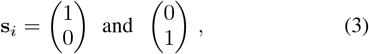

respectively. The accumulated payoff, *U*_*i*_, of player *i* is thus given by

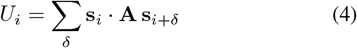

where **A** denotes the payoff matrix, Eq. 1, and the sum runs over the four neighbouring sites *i* + δ.

### C. Competition

Individuals compete for reproduction by comparing their payoffs to those of their neighbours. Again in the simplest case a focal player compares their payoff to a randomly chosen neighbour. If the neighbour performs better than the focal, they probabilistically adopt the neighbour’s strategy (cultural evolution) or, equivalently, the focal is replaced by clonal offspring of the neighbour (genetic evolution). The strategy distribution in the population then evolves by repeatedly sampling a focal player *i* uniformly at random who compares their payoff *U*_*i*_ to one of their randomly chosen neighbours *j* and adopts player *j*’s strategy with probability

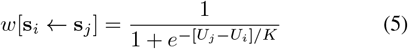

where *K* quantifies the noise in the updating process [27].

### D. Neighbourhoods & sub-populations

Traditionally the interaction and competition neighbourhoods are assumed to be identical. Indeed, fully overlapping neighbourhoods are best for cooperation [28, 29]. Similarly, clustering provides stronger support for cooperation in smaller neighbourhoods [25, 30]. Complementing the traditional, overlapping case we consider the opposite extreme where the two neighbourhoods are entirely disjoint.

In both cases individuals interact with their *k* = 4 nearest neighbours to the north, east, south and west (von Neumann neighbourhood). In the traditional scenario all players also compete with the same set of neighbours. Whereas, in the disjoint scenario, players still compete with same number of neighbours, *l* = 4, but from a disjoint set, given by their *second-nearest*, diagonal neighbours (to the north-east, south-east, south-west, and north-west), see Fig. 1. Thus, competing individuals never directly interact but share interaction partners. Incidentally, all second-nearest neighbours again form a lattice, or, more precisely, two disjoint sub-lattices. Effectively, this divides the population into two distinct sub-populations with interactions exclusively *between* sub-populations but competition exclusively *within* each sub-population. The resulting structure is closely related to models for studying mutualistic interactions between species [32, 33].

**FIG. 1.**
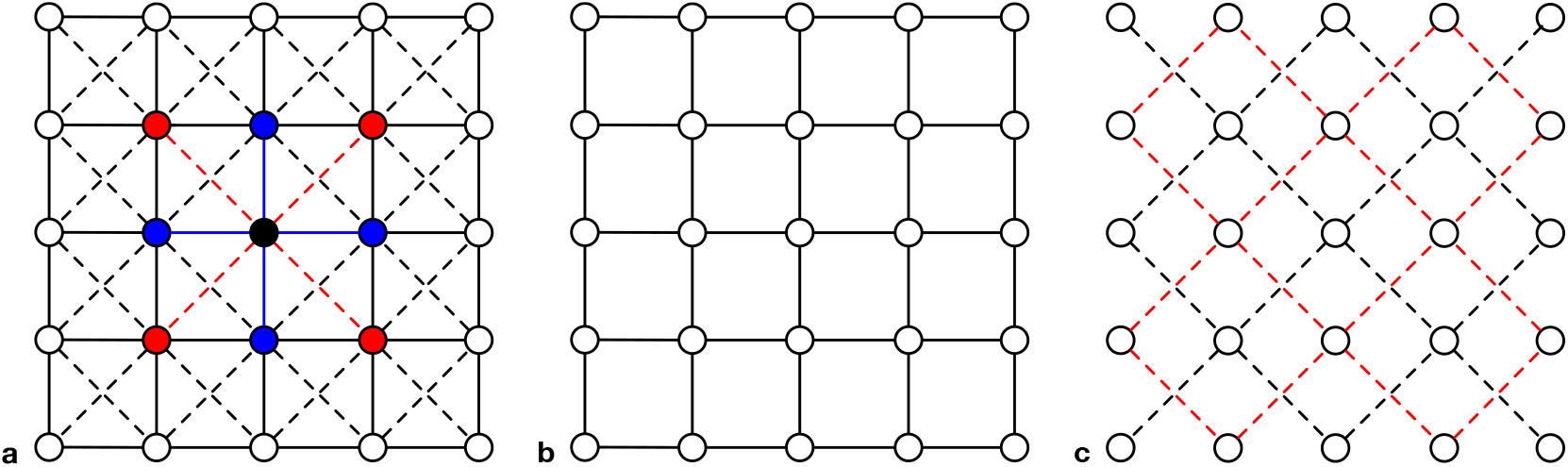
(colour online) **a** Schematic depiction of the disjoint interaction (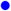, solid lines) and competition (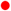, dashed lines) neighbourhoods of the focal individual (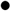) on a square lattice. **b** The interaction network is the standard 2D square lattice with degree *k* = 4 and includes all members of the population. **c** In contrast, the reproduction (or imitation) network connects the *l* = 4 second-nearest neighbours along the diagonals. This results in two disjoint sub-lattices (black and red dashed lines, respectively) of equal sizes, provided that *N* is even on an *N × N* lattice. For interactive online simulations of the donation game see [31].

### E. Methods

The macroscopic state of the population is characterized by frequency *ρ*(*t*) of cooperators in the traditional setup and *ρ*_1_(*t*) and *ρ*_2_(*t*) for the frequency of cooperators in each sub-lattice, respectively. Evolution stops once the population has reached an absorbing state. The traditional scenario admits two absorbing states with *ρ*(*t*) = 0 (extinction of cooperators) and *ρ*(*t*) = 1 (fixation of cooperators). In contrast, the disjoint setup admits four absorbing states with *ρ*_*a*_(*t*) = 0 or 1, such that cooperators vanish or fixate in each sub-lattice (*a* = 1 or 2). In the long run the population inevitably reaches one of those absorbing states. More importantly, however, the population can spend exceedingly long times in quasi-stationary states, which can be characterized by the average frequency of cooperators 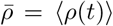 and 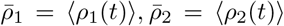, respectively.

The average frequencies are determined by Monte Carlo (MC) simulations as a function of the cost-to-benefit ratio *r* and noise *K* (temperature) by averaging over a sampling time *t*_*s*_ after a relaxation time *t*_*r*_. The time is measured in number of Monte Carlo steps (MCS). In a population of size *N* each MCS represents *N* elementary updating steps, such that every player has, on average, one chance to propagate their strategy, which is also termed one generation. Naturally, these measurements are conditioned on not reaching any absorbing state, which necessitates sufficiently large lattice sizes to prevent accidental extinction of one type. The linear lattice size *L*, as well as the values of *t*_*r*_, *t*_*s*_ are adjusted to the dynamical behaviour depending on *r* and *K*.

The runtime of MC simulations can be significantly reduced for cases where the frequency of the sparse strategy is very small. Naturally this is particularly important to understand and reveal the details of the transitions to extinction of either type. In this case all *n*(*t*) players with the sparse strategy at time *t* are labeled 0, …, *n*(*t*) − 1. The selection of candidates for updating is limited to those sites as well as their competing neighbours. After every successful imitation (replication) event *n*(*t*) changes by ±1 and the labels are updated accordingly. This method is much less affected by the lattice size and allows to consider populations that require relaxation and sampling times of *t*_*r*_, *t*_*s*_ *>* 10^7^ MCS. Using this method we are able to confirm the power law scaling of frequencies when approaching extinction.

An equally useful macroscopic measure to characterize the dynamical features are the fluctuations of the frequency of cooperation, *χ*. While relevant to characterize phase transitions between distinct dynamical regimes in either scenario, we refer to [26] for the traditional case and use it here only for the sub-lattices, given by

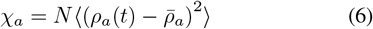

where *a* = 1, 2 refers to one or the other sub-lattice and ⟨…⟩ denotes the average over the sampling time. In the limit *N* → ∞, *χ*_*a*_ becomes constant [34] and *χ*_*a*_ = 0 holds for homogeneous sub-lattices.

The MC simulations are complemented by results based on pair approximation and the four-site dynamical cluster method. The latter is a straightforward extension of the meanfield and pair approximations. Interestingly, pair approximation fails to capture some qualitative effects observed in the simulations. Those require the increased accuracy of larger unit clusters. The technical details of these approximations are given in the literature of non-equilibrium statistical physics [35, 36] and spatial evolutionary games [11, 37]. In essence, the four-site approximation describes the state based on configuration probabilities of 2 × 2 clusters of individuals and the microscopic transitions between them. The number of independent variables and transitions can be significantly reduced by symmetries and compatibility conditions. This yields a set of differential equations that translate the effects of the elementary imitation processes on the time-dependent configuration probabilities of the 2 × 2 clusters into global frequencies of cooperation in the two sub-lattices. This approximation turns out to be sufficient to qualitatively reproduce the MC results.

## II. OVERLAPPING NEIGHBOURHOODS

Let us first consider the traditional case with identical interaction and competition neighbourhoods. This complements and confirms previous results for the weak prisoner’s dilemma [11, 66] and extends them to the donation game. Moreover, this provides a valuable basis for later comparisons to the dynamics with disjoint neighbourhoods. More specifically, we consider the average frequency of cooperation in the donation game as a function of the cost-to-benefit ratio *r* at a fixed noise level, *K* = 0.1. In well-mixed populations defection dominates for *r >* 0, while for negative costs, *r <* 0, the interaction turns into a harmony game where cooperation is the dominant strategy. In spatial games, cooperators and defectors co-exist in the vicinity of *r* = 0. Results from MC simulations, pair- and four-site approximations are depicted in Fig. 2. Typical spatial configurations of cooperators and defectors near their respective extinction threshold are shown in Fig. 3. For *r >* 0 small, isolated clusters of cooperators persist and withstand the threat of exploitation by defectors. In contrast, for *r <* 0 mostly isolated or pairs of defectors persist through spiteful actions that harm their neighbours more than themselves.

**FIG. 2.**
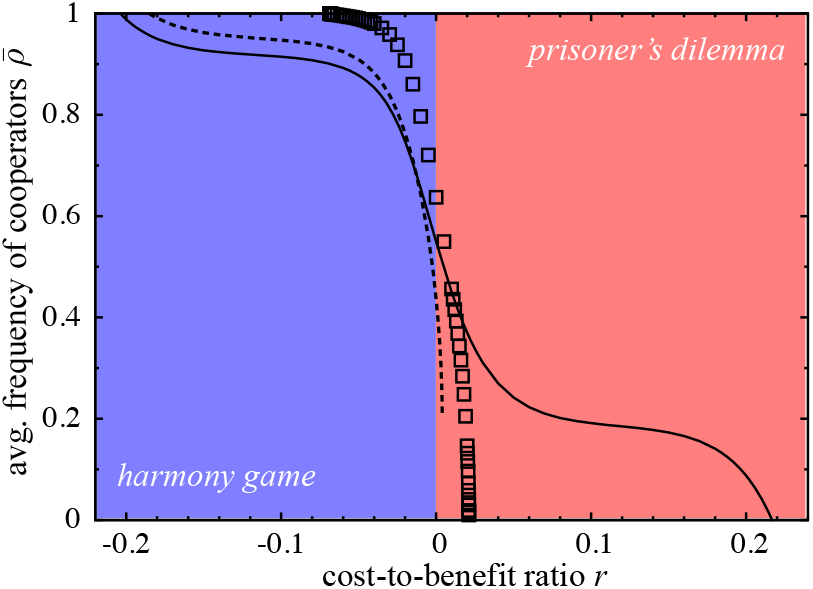
(colour online) Average frequency of cooperators, 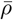, as a function of *r* with overlapping interaction and competition neighbourhood for noise level *K* = 0.1 for MC simulations (□), pair- (solid line) and four-site (dashed line) approximations.

**FIG. 3.**
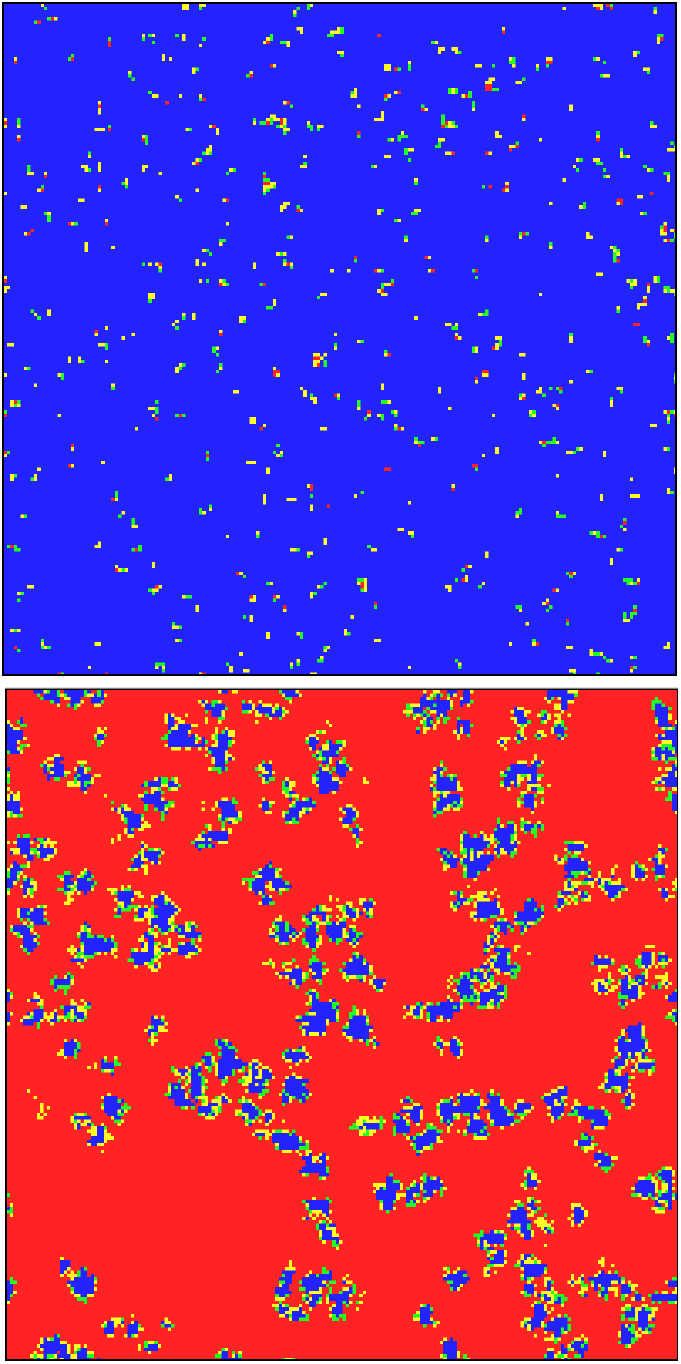
(colour online) **a** defectors (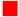and 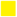) survive through spite in the harmony game, *r* = *−*0.04. **b** clustering enables cooperators (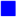 and 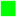) to persist in the donation game, *r* = 0.02. Individuals that updated their strategy in the last MC step (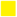 and 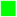) visualize the activity in the population. The noise in both panels is *K* = 0.1. For interactive online tutorials see [31].

### A. Phase transitions: extinction of cooperators and defectors

Spiteful defectors survive with noise-dependent (and sometimes extremely low) frequencies. In general, the frequency of cooperators decreases monotonously for increasing *r*. Cooperators and defectors can only co-exists for *r*_*D*_ *< r < r*_*C*_ with *r*_*C*_ = 0.021105(2) and *r*_*D*_ = −0.068768(2) based on MC results obtained for *K* = 0.1. Pair approximation significantly overestimates the range of coexistence and the four-site approximation predicts a first order transition at *r* = 0.003892 from 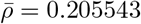 to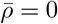.

In contrast, simulation results show that the extinction of cooperators as well as defectors are consistent with theoretical predictions of non-equilibrium critical phase transitions [38, 39] that fall into the directed percolation universality class [40, 41]. In two-dimensional systems this transition is characterized by a power law decrease in the frequency of the vanishing strategy, 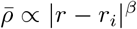 where *i* ∈ *C, D* with *β* = 0.58 and *r* → *r*_*i*_ from below for *i* = *C* but from above for *i* = *D* [11, 26]. Critical transitions are accompanied not only by diverging fluctuations, but also diverging spatial and temporal correlation lengths. Thus, accurate MC simulations in the vicinity of critical transitions require not only large lattices, *L >* 1000, to prevent accidental extinction but also long relaxation and sampling times, *t*_*s*_, *t*_*r*_ *>* 10^6^ MCS, for representative measurements.

The power law decrease of defectors is not visible in Fig. 2 because of the extremely low frequencies of 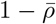 when *r* → *r*_*D*_ from above. However, the similarity of the extinctions of cooperators and defectors becomes evident in Fig. 4.

**FIG. 4.**
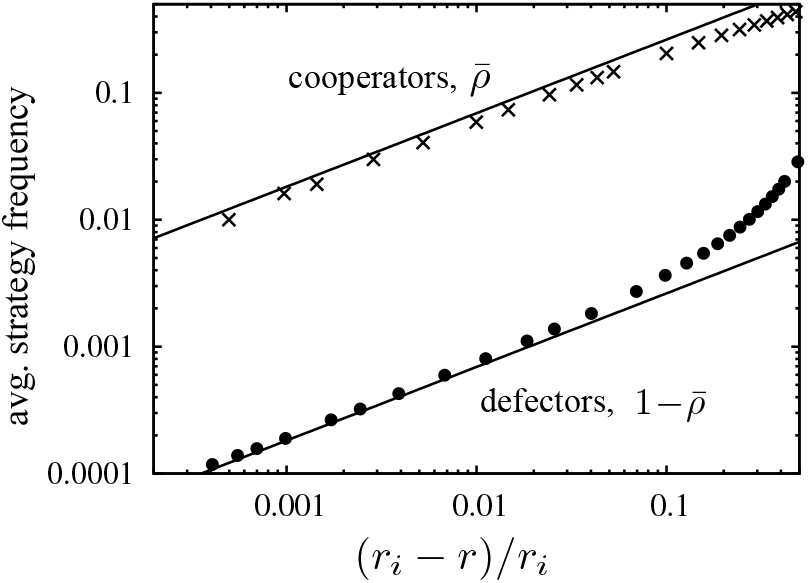
The average frequency of cooperators, 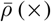, and defectors, 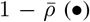, scales with a power law when approaching the critical cost-to-benefit ratios, *r*_*C*_ and *r*_*D*_ from below and above, respectively (*K* = 0.1). The theoretical expectation is a power law with *β* = 0.58 (solid line).

The extinction of defectors can take exceedingly long times because, for negative *r*, solitary defectors do well in spatial interactions by harming their neighbours more than them-selves. As a consequence, cooperating neighbours likely imitate spiteful defection, but then both suffer from mutual defection. Subsequently, either one switches back to cooperation and again an isolated defector remains. This process effectively resembles a branching-coalescing-annihilating random walk [42, 43]. On large lattices the coalescence time increases. Spontaneous annihilation of solitary defection is allowed by Eq. 5 but with a probability approaching zero in the low noise limit. The same applies to branching events. Extremely small fractions of defectors, 1 − *ρ* ≃ 0.00001, persist for a long time, *t >* 10^7^ MCS in simulations for *L* = 10^4^, *K* = 0.1 and *r* = −0.07. In fact, for such low noise the probability to eliminate the last defector is merely about *e*^*−*(1+4*r*)*/K*^ per MCS (or less than 3 · 10^*−*4^% for the above values).

### B. Effects of noise

The extinction threshold of cooperators, *r*_*C*_(*K*), as well as defectors, *r*_*D*_(*K*), depends on the noise *K*. The slow coalescence and the unlikely elimination of the last defector both cause technical difficulties to determine *r*_*D*_(*K*) at lower noise levels. Fig. 5 illustrates the persistence of defectors at very low frequencies with little noise, *K* = 0.06, and *r* approaching *r*_*D*_(0.06) ≃ −0.08615(5). Also note that for these parameters the stationary states are reached only after long relaxation times of *t*_*r*_ *>* 10^7^ MCS. Those results are effectively inaccessible without the labelling technique of the rare strategy as outlined above. The time series are consistent with the general features of directed percolation universality class, which predicts diverging relaxation time and fluctuations when *r* → *r*_*D*_ in the limit *L* → ∞.

**FIG. 5.**
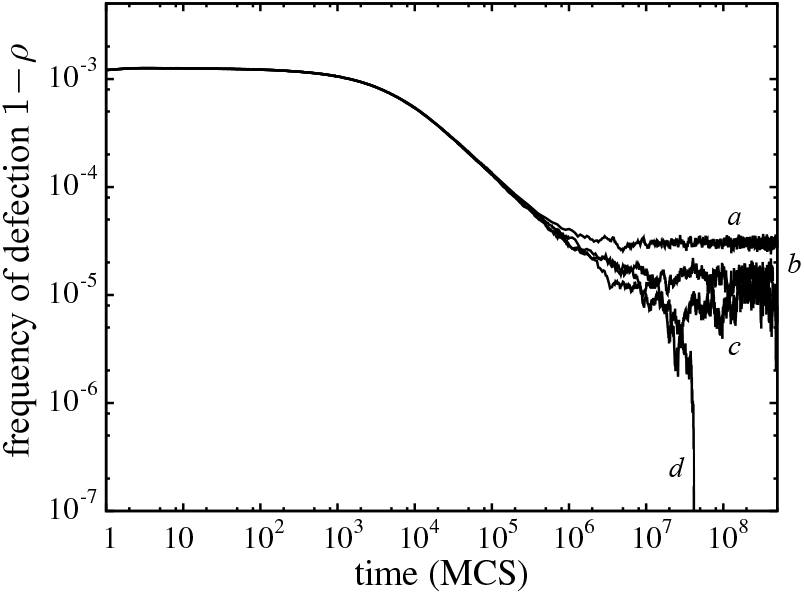
Defector frequencies over time for spiteful behaviour in the spatial donation game on an *L × L* lattice with overlapping interaction and competition neighbourhoods at low noise, *K* = 0.06. Parameters: *L* = 10000 and (a) *r* = *−*0.0856, (b) *−*0.0860, (c) *−*0.0861, and (d) *−*0.0862.

Most of the literature on spatial donation games focussed on fixed noise levels, *K*. Here we extend the analysis to explore the effects of noise and repeat the above simulations for different *K* to determine the region of co-existence, *r*_*D*_(*K*) *< r < r*_*C*_(*K*), as a function of *r* and *K*, see Fig. 6. The results illustrate that spiteful behaviour is most significant at low *K* and then decreases with increasing *K*. Most importantly, however, Fig. 6 reveals the existence of an optimal noise level, *K* ≃ 0.3, where *r*_*C*_(*K*) exhibits a maximum. In the high noise limit, *K* → ∞ the extinction thresholds converge to *r*_*C*_(*K*), *r*_*D*_(*K*) → 0 and for *K* → 0, *r*_*C*_(*K*) 0 holds. Although, for other update rules *r*_*C*_(0) *>* 0 may hold [44].

**FIG. 6.**
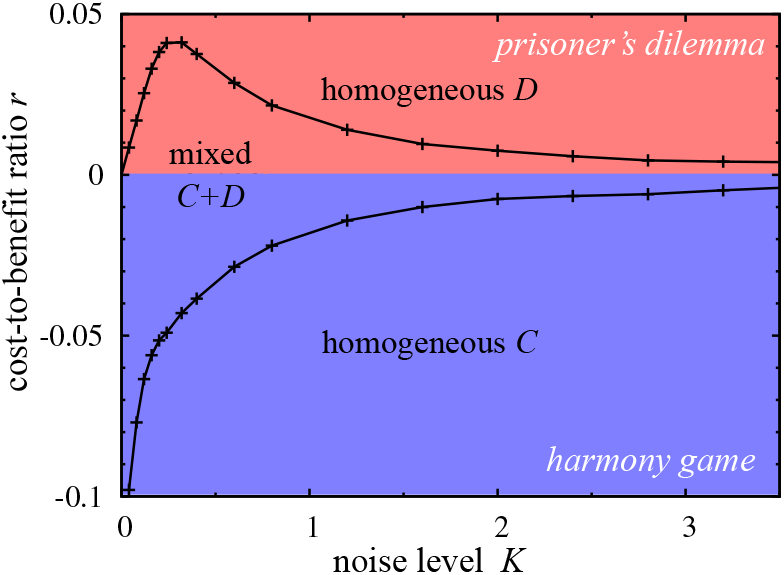
(colour online) The stationary state as a function of *K* and *r* in the donation game on square lattices with overlapping interaction and competition neighbourhoods. Symbols depict the values of *r*_*C*_ (*K*) and *r*_*D*_ (*K*) as derived from MC simulations.

In the high noise limit, *K* → ∞, Eq. 5 becomes a coin toss and our model becomes equivalent to the voter model [42, 45]. In this limit the domain growth process is extremely slow (and depends on the spatial dimension). More precisely, on square lattices the average size of homogeneous domains increases with log *t* [46].

## III. DISJOINT NEIGHBOURHOODS - TWO SUB-POPULATIONS

In the second scenario, interactions again happen with the *k* = 4 nearest neighbours. However, competition does not happen among the same set of neighbours but an entirely disjoint set of the same size given by the *l* = 4 second nearest neighbours (along the diagonals). The updating of strategies also follows Eq. 5, as before.

In a social context this describes situations when players adopt strategies not from their interaction partners but from those of their neighbours. Incidentally this results in two disjoint competition lattices (see Fig. 1). This type of competition is beneficial for players in several spatial games. For example, the hawk-dove or snowdrift games [23] admit two pure Nash equilibria where one player cooperates and the other defects. Those equilibria cannot be realized with a single competition lattice but are feasible with two disjoint sub-lattices.

More specifically, the disjoint scenario admits four absorbing states because strategies can fixate independently on each sub-lattice: *CC, CD, DC*, and *DD*. For sub-lattices that are composed of different strategic types, the configuration looks like a checkerboard. Most intriguingly, such asymmetric configurations between the sub-lattices can arise spontaneously on an ephemeral, local scale of (almost) frozen regions or in the form of globally absorbing states with homogenous cooperation or defection in either sub-lattice.

### A. Phase transitions

The frequency of cooperation in each sub-lattice as a function of *r* is shown in Fig. 7. The noise is fixed at *K* = 0.5, which is close to the optimal value for supporting cooperation in the traditional scenario, c.f. Fig. 6. Interestingly and in contrast to the traditional scenario it turns out that spiteful behaviour is no longer feasible and thus we limit the analysis to *r >* 0.

**FIG. 7.**
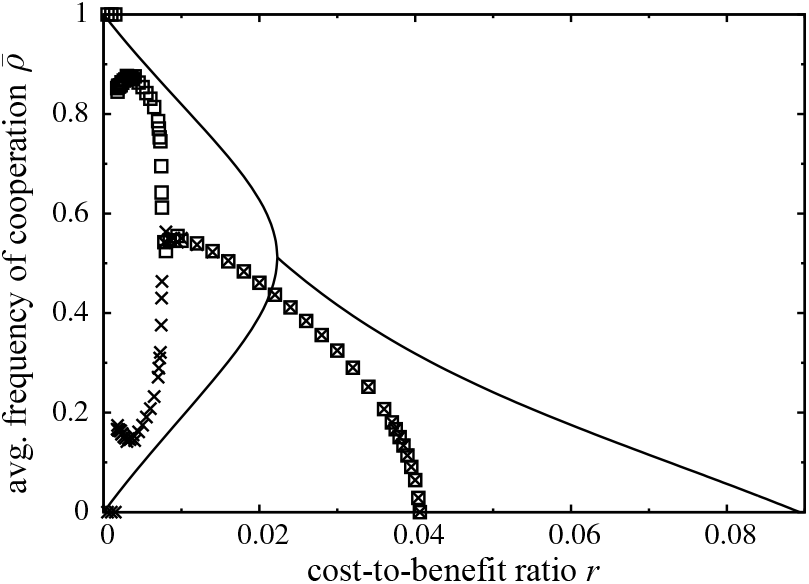
Average frequency of cooperation, 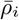, as a function of *r* for *K* = 0.5 in each of the two sub-lattices, *i* = 1, 2. The results from MC simulations (□ and *×* symbols for each sub-lattice) are complemented by predictions of the four-site dynamical cluster method (solid lines).

Figure 7 depicts four dynamical regimes separated by three critical phase transitions: *(i)* for large *r* clustering benefits are insufficient to sustain cooperation; *(ii)* for *r*_*s*_ *< r < r*_*C*_ cooperators persist in a symmetric phase with equal frequencies in both sub-lattices; *(iii)* for *r*_*a*_ *< r < r*_*s*_ cooperators persist in an asymmetric phase with complementary frequencies in the sub-lattices; and *(iv)* for *r < r*_*a*_ the populations ends up in one of the fully asymmetric absorbing states *CD* or *DC*.

The critical phase transitions are accompanied, and indeed defined by diverging fluctuations when approaching the critical points, see Fig. 8. Unsurprisingly, the fluctuations in each sub-lattice are the same, χ_1_ = χ_2_, in the symmetric phase,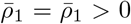. In the polarized state of the anti-ferromagnetic Ising model ρ_1_ = 1 − ρ_2_ holds for spins pointing up or down, respectively. However, in the asymmetric phase this equality holds only approximately between the two sub-lattices and gives rise to small but systematic differences in the fluctuations. The scaling of the divergence of χ_*i*_ confirms the characteristics of the directed percolation and Ising-type universality classes, respectively.

**FIG. 8.**
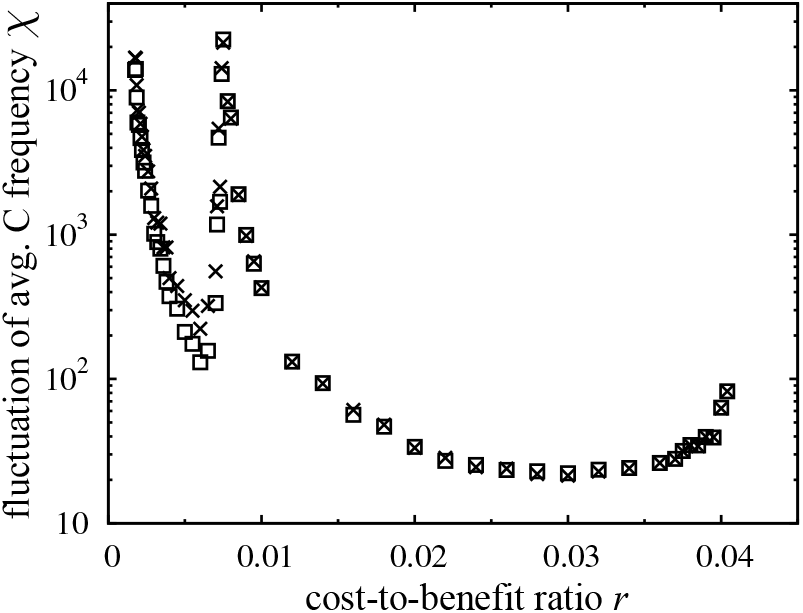
Simulation results for fluctuations of the cooperator frequencies, *χ*_*i*_, in each sub-lattice, *i* = 1, 2 (□ and *×*). Parameters: see Fig. 7.

Based on MC simulations with *L >* 1000, and *t*_*r*_, *t*_*s*_ ≥ 10^5^ MCS, cooperators go extinct in both sub-lattices simultaneously for *r* → *r*_*C*_ = 0.04055(2) from below. This critical phase transition is similar to the traditional scenario and matches theoretical expectations for the directed percolation universality class in 2D, which predicts a power law 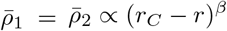 with an exponent *β* = 0.58.

### B. Symmetry breaking

The most striking feature is the appearance of spontaneous symmetry breaking between the levels of cooperation in the two sub-lattices 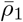 and 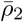 if *r < r*_*s*_. The MC data for this second critical phase transition at *r*_*s*_ are consistent with the theoretical expectation for the Ising-type universality class, which predicts 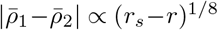 in the limit *r* → *r*_*s*_ from below with *r*_*s*_ = 0.00741(5). In fact, there are two equivalent spatial arrangements of strategies reminiscent of the twofold degeneracy of the ordered phases in the anti-ferromagnetic Ising model in the absence of an external magnetic field [47–49]. In both cases the ordered regions admit two equivalent configurations by exchanging the strategic types between sub-lattices, or by flipping spin orientations, respectively. The imitation of second neighbours on a square lattice support the formation of large domains of such ordered configurations. This amounts to spontaneous symmetry breaking between the two sub-lattices, such that in one sub-lattice cooperators are more frequent but defectors in the other.

Typical spatial configurations of cooperators and defectors in the symmetric, *r > r*_*s*_, and asymmetric *r < r*_*s*_, phases are shown in Fig. 9. The symmetry breaking is accompanied by a power-law divergence in the relaxation times, *t*_*r*_. This is also referred to as critical slowing down on both sides of the critical point [34, 50, 51]. This general and universal feature depends on the spatial dimension and may be related to the vanishing force (at the critical point) driving the formation of the stable phase, regardless of whether the symmetric or asymmetric one.

**FIG. 9.**
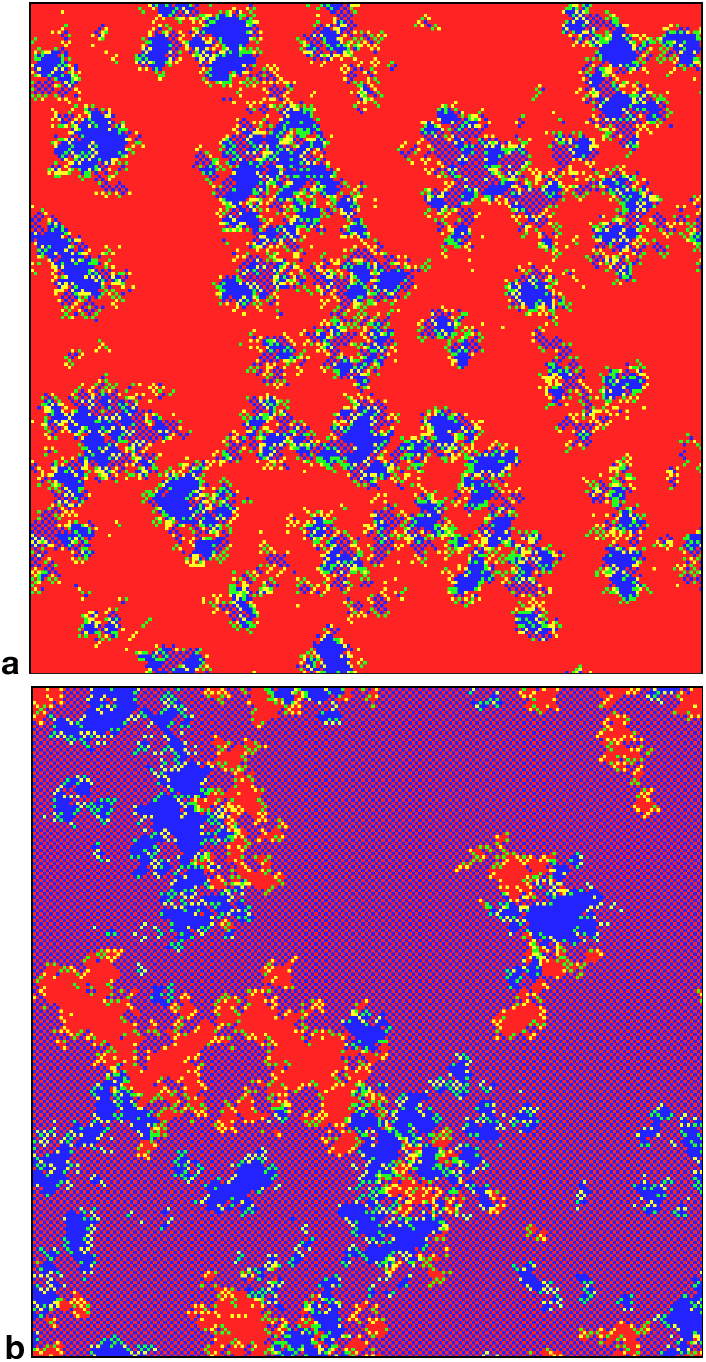
(colour online) Typical spatial configuration of cooperators (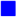and 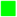) and defectors (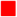 and 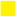) in **a** the symmetric phase with equal frequencies in both sub-lattices with *r* = 0.035, and **b** the asymmetric phase essentially complementary frequencies in each sub-lattice with *r* = 0.005. Individuals that updated their strategy in the last MC step (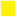and 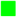) visualize the activity in the population. The noise in both panels is *K* = 0.5. For interactive online tutorials see [31].

Similar spontaneous symmetry breaking in spatial evolutionary game theory models has been reported for interactions with *multiple* layers [33, 52–55].

### C. Asymmetric dynamics

Symmetry breaking results in different average payoffs for players on the two sub-lattices. For example, if 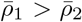 then players in sub-lattice 2 (with less cooperation) have, on average, higher payoffs than those in 1. This can be interpreted as one sub-population exploits the other, at least on average. Since the sub-lattices are equivalent, the opposite situation 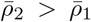 can also occur with a probability that depends on the initial configuration as well as the noise in the evolutionary process.

The existence of two equivalent stable phases, *CD* and *DC*, results in significant delays in the relaxation process. This is particularly pronounced when starting from a symmetrical initial state. In this case both phases can occur separately by forming domains with an increasing average size 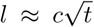, where *c* decreases monotonously with the noise level. These types of domain growing processes are well investigated for the kinetic Ising model [56, 57]. Macroscopic variations in the asymmetry, 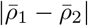, (magnetization) arise only once the average linear size of the domains, *l*, becomes comparable to the system size *L*.

This delay in reaching a relaxed equilibrium state becomes apparent in the time series of *ρ*_1_(*t*) and *ρ*_2_(*t*) for different initial configurations, see Fig. 10. Because two equivalent, complementary asymmetric states exist, symmetries in the initial configuration cause delays in the convergence towards one or the other. Fortunately, these delays can be easily and efficiently avoided by using initial configuration with asymmetric frequencies.

**FIG. 10.**
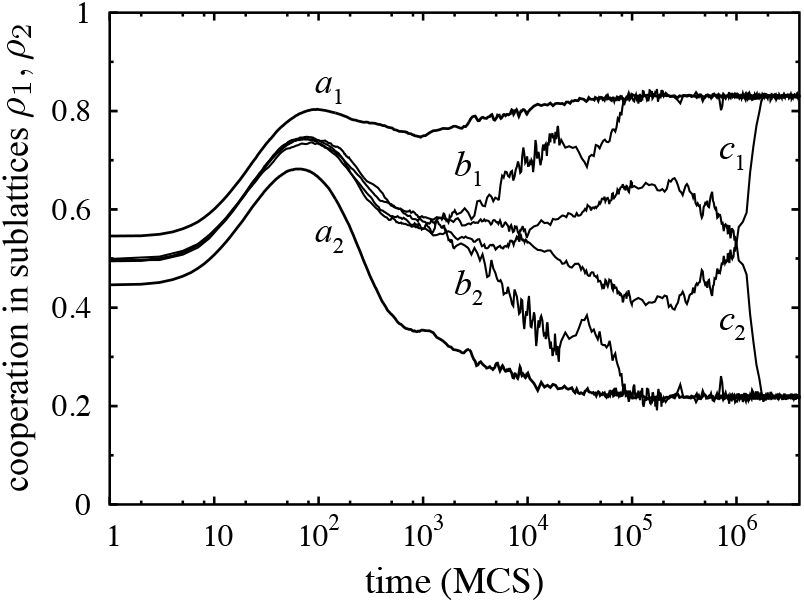
Time series of the frequency of cooperation each sub-lattice in the region of spontaneous symmetry breaking for *K* = 0.5 and *r* = 0.002. The asymmetric equilibrium is reached quickly for an asymmetric initial configuration (lines *a*_1_, *a*_2_) for *L* = 2000. In contrast, for symmetrical initial frequencies the relaxation takes significantly longer even for smaller lattices (*b*_1_, *b*_2_ with *L* = 600 and *c*_1_, *c*_2_ with *L* = 1200) due to the slow domain growing process.

#### Cluster approximation

The dynamical features and symmetry breaking, in particular, are qualitatively reproduced by the four-site dynamical cluster approximation (solid lines in Fig. 7). Notable differences are that the frequency of co-operation vanishes linearly, 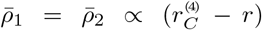 for 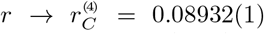 from below. The superscript ^(4)^ refers to results based on the four-site approximation. The spontaneous symmetry breaking exhibits parabolic behaviour, 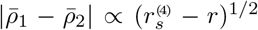, for 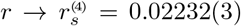 from below. These systematic deviations reflect the relevance of long-range correlations that cannot be captured in this approximation. In order to determine the exponents of the different power laws, it is necessary to integrate the dynamical equations for very long times (*t*_*r*_ *>* 10^4^) in the vicinity of the extinction of cooperators, 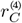, and the symmetry breaking transition, 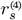. The trajectories converge monotonously (or, more precisely, exponentially) to the stationary state.

### D. Absorption into fully asymmetric states

The asymmetry in cooperation between the sub-lattices, 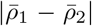, has a maximum at *r* ≃ 0.003. For smaller *r* the fluctuations in the strategy frequencies, *χ*_*i*_, keep increasing but result in a decrease in the asymmetry. Except for very small 0 *< r <* 0.0018, where the population tends to get absorbed in one of the fully asymmetric states, *CD* and *DC*, or homogeneous defection, *DD*, through a complex process dependent on the initial state and the stochastic elementary steps.

In finite systems, the fully asymmetric end states occur for higher values of *r* due to fluctuations. As illustrated in Fig. 11, in one simulation the early extinction of cooperators in one sub-lattice is followed by a decrease in the other, resulting in homogeneous defection, *DD*. The other simulation depicts the trajectory when, after huge, correlated fluctuations in both sub-lattices, evolution ends in the fully asymmetric *CD* state. As soon as one sub-lattice becomes homogeneous, regardless of whether *C* or *D* fixated, defectors in the other sub-lattice have a systematic advantage. Nevertheless, on small lattices, all four absorbing states can be reached due to the stochastic nature of the evolutionary process.

**FIG. 11.**
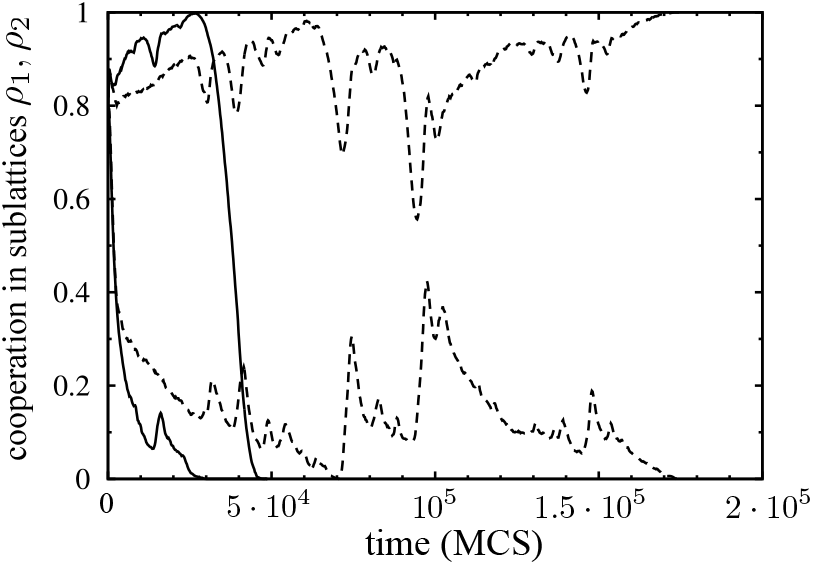
Time series of cooperation frequencies, 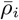, in the two sublattices, *i* = 1, 2, for weakly perturbed initial states with *ρ*_2_(*t* = 0) − ρ_1_(*t* = 0) = 0.05 and *r* = 0.0018, *K* = 0.5, on an 1000 *×* 1000 lattice. One trajectory ends in homogeneous defection (solid line) and the other in the fully asymmetric *CD* state (dashed line).

Bursts may cause dramatic variations in the spatio-temporal patterns. The dynamics on small lattices is easily driven into one of the absorbing states (see Fig. 11). On large lattices similar extinctions can occur within limited regions and followed by a slow recovery, see Fig. 12. The large bursts and the subsequent slow relaxation processes cause serious technical difficulties in the accurate determination of the average frequencies of cooperation. For example, the highest values of χ in Fig. 8 are obtained from MC simulations on a large lattice, *L* = 4000, with *t*_*r*_ = 10^6^ MCS and *t*_*s*_ *>* 5 10^7^ MCS.

**FIG. 12.**
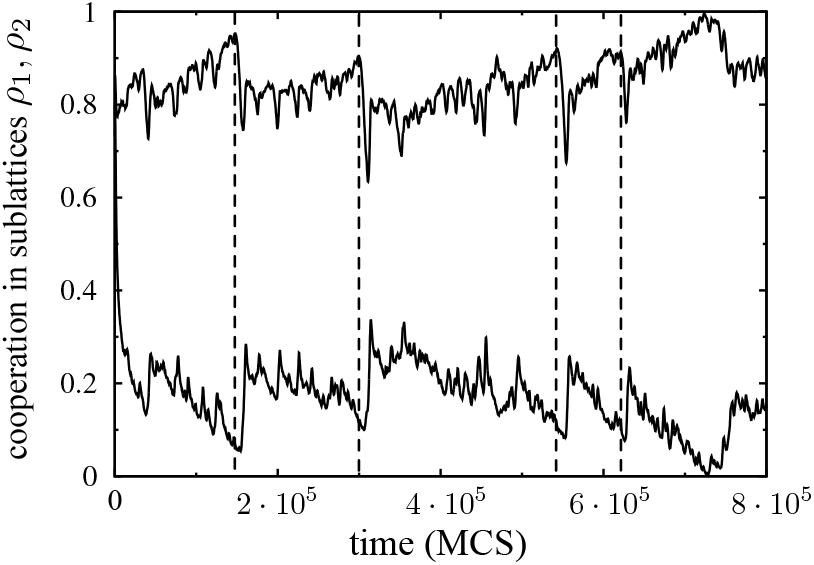
Time series of cooperation frequencies as in Fig. 11 but on a large 2000 *×* 2000 lattice. Crashes in cooperation in the sub-lattice with more cooperators are followed by a delayed increase in cooperation in the other sub-lattice (the dashed vertical lines highlight the onset of a few of those crashes).

Local maxima of *ρ*_2_(*t*) are often followed by a rapid and huge decrease in cooperation with a slow recovery. With some delay the variation is mirrored in *ρ*_1_(*t*). Similar phenomena are observed in a two-layer evolutionary donation game [33]. In both systems, the emergence of complex self-organizing, microscopic patterns control the macroscopic fluctuations. In the asymmetric phase, *CD* islands are located in a sea of *DC*. Along the boundary, small *CC* and *DD* islands are created by the stochastic strategy updates. *DD* islands can move, grow, shrink, and vanish at random. However, if a *DD* islands exceeds a critical size then it expands inside the *DC* domain and decreases *ρ*_2_(*t*). This extension of the *DD* domain is blocked and reversed when colliding with a small *CC* cluster. The invasion of *CC* into large *DD* domains becomes observable macroscopically as simultaneous increases in *ρ*_1_(*t*) and *ρ*_2_(*t*). Finally, *CC* domains are prone to invasion by either *CD* or *DC*, which restores the original asymmetry (or its opposite). This cyclical invasion drives the macroscopic fluctuations in *ρ*_1_(*t*), *ρ*_2_(*t*). The *DD* bursts become larger and rarer for decreasing *r* and result in diverging fluctuations (see Fig. 8), which eventually drive the population into one of the fully asymmetric absorbing states, *CD* or *DC* for *r < r*_*a*_ = 0.0018.

In fact, for small *r* the fully asymmetric absorbing states are the *preferred* outcomes because once one sub-lattice has become homogeneous, the payoffs of those individuals no longer matter and the dynamics on the other sub-lattice is dominated by random drift: Eq. 5 essentially reduces to a slightly biased coin-toss for small payoff differences. Regardless of whether *C* or *D* fixated in one sub-lattice, the difference in payoffs for cooperators and defectors in the other lattice is 4*r*, which reflects the small bias in favour of defectors. Thus, the fixation probability of each strategic type on the other sub-lattice is essentially given by their frequency at the time when the first sub-lattice became homogenous. For example, if a sub-lattice becomes homogeneous in *C* then the frequency of *C*’s in the other sub-lattice is likely *<* 1*/*2, due to the asymmetry. Thus, the *CD* absorbing state is more likely than *CC*. Conversely, if one sub-lattice becomes homogeneous in *D*, then the frequency of *C*’s in the other sub-lattice is likely *>* 1*/*2 and hence the *DC* absorbing state is more likely than *DD*. Finally, it is more likely that one layer becomes homogeneous in *D* because isolated *C*’s (or small *C* clusters) are more easily eliminated than *D*’s. Thus, even though all four absorbing states may be reached, in principle, the probabilities are very different and most likely approach heterogeneous absorbing states, leaving homogeneous cooperation across lattices the least likely outcome.

Burst dynamics has been reported and discussed in some other systems [58], including evolutionary games [33, 59]. In the present model, however, bursts result from the combination of cyclic dominance in the macroscopic invasion rates and the nucleation processes [57, 60, 61]. Similar phenomena are described in a simple evolutionary game where the three-state Potts model is extended by a rock-paper-scissors type interaction [62].

#### Cluster approximation

Interestingly, the divergence of relaxation times also affects the dynamical cluster method. In the limit *r* → 0, where MC simulations are subject to bursts, the convergence fundamentally changes and happens through slowly damped oscillations as visualized by spiralling trajectories in the *ρ*_1_(*t*)−*ρ*_2_(*t*) plane, see Fig. 13. The damping decreases with *r* and hence convergence becomes slower. Also note that the trajectory gets extremely close to the two axes, which mark extinction of cooperators (or defectors). More specifically, *ρ*_1_(*t*) and 1 − *ρ*_2_(*t*) become smaller than 10^*−*6^ along the plotted trajectory. This indicates that in any finite system the dynamics likely ends up in an absorbing state as a consequence of stochastic fluctuations in the frequency of cooperators or defectors on either sub-lattice.

**FIG. 13.**
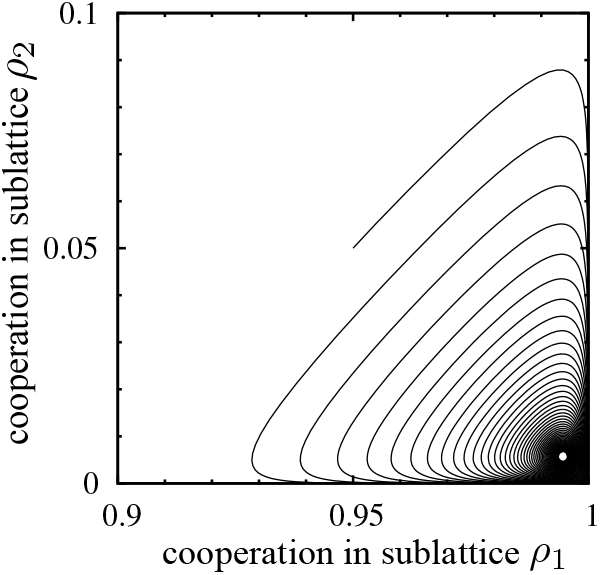
Spiralling trajectory in the *ρ*_1_(*t*) − ρ_2_(*t*) plane for the dynamical equations of the four-site cluster method with *K* = 0.1 and *r* = 0.0001. The integration is stopped after *t* = 10^4^ and ends close to the stationary state at the centre of the damped oscillations.

### E. Effects of noise

For a more complete picture on the effects of noise, the simulations are repeated for fixed *r* = 0.005 and varying *K*, see Fig. 14. Interestingly, cooperators vanish in the limit *K* → 0 as well as at high noise levels. For an intermediate range of *K* spontaneous symmetry breaking occurs. More specifically, note that asymmetries arise once the total average frequency of cooperation becomes sufficiently high, approximately 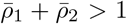. The asymmetric phase is bounded by symmetrical phases for both higher and lower *K*. Interestingly, the dynamical four-site cluster approximation is capable of qualitatively capturing all these dynamical features.

**FIG. 14.**
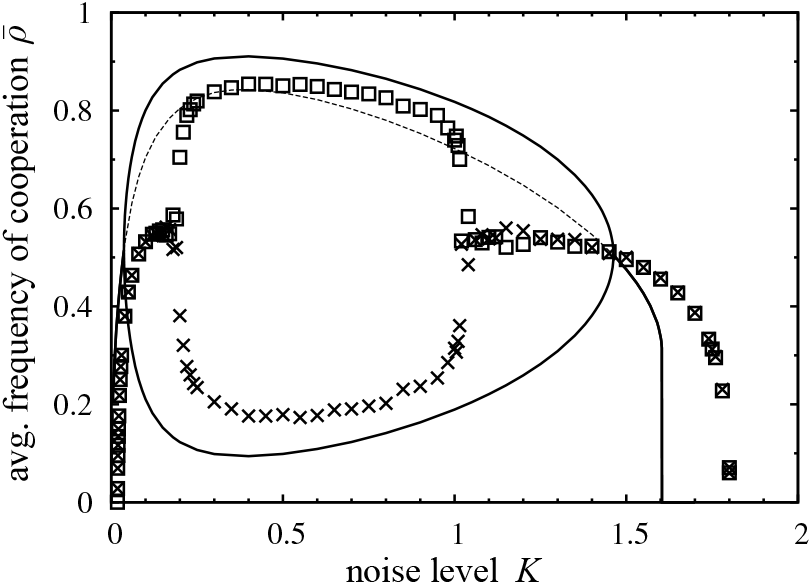
The average frequency of cooperation, 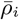, in the two sublattices, *i* = 1, 2, for *r* = 0.005 as a function of the noise *K*. Simulation results are shown for each sub-lattice (□ and *×*) and complemented by predictions of the four-site dynamical cluster method (solid lines), including the unstable symmetrical solution (dotted line).

Again, in the high noise limit, *K* → ∞, our model becomes equivalent to the voter model. More specifically, for the disjoint neighbourhoods, the dynamics on the two sub-lattices become decoupled and the spatial dynamics corresponds to two independent voter models on each sub-lattice. Similar behaviour is expected for any *r* after one sub-lattice reached a homogeneous, absorbing state. The payoffs to cooperators and defectors in the other sub-lattice are identical and reduce to a voter model on the heterogeneous sub-lattice.

## IV. SUMMARY & DISCUSSION

Limited local interactions in spatially structured populations have long been recognized as a potent promoter of co-operation [10, 25]. Here we report surprisingly different dynamics for the evolutionary donation game, depending on whether individuals compete with the neighbours they also interact with – as is traditionally the case – or compete with those two steps removed. More generally, the degree to which interaction and competition neighbourhoods overlap affects the fate of cooperation and gets more challenging with larger competition neighbourhoods or less overlap between the two [29, 30, 63, 64].

A particularly interesting and simple setup is given by a square lattice with four neighbours because the number of first and second neighbours (along the diagonal) is the same. However, the set of diagonal neighbours actually forms two disjoint sub-lattices (c.f. Fig. 1) and effectively splits the population into two sub-populations. An important consequence is that, instead of the two absorbing states of all cooperation or all defection, the population now admits four absorbing states with all combinations of homogeneous cooperation or defection in one or the other sub-lattice.

Whether competition arises on a single lattice or two disjoint sub-lattices has essentially no impact on some aspects of the dynamics. In particular, the characteristics of the extinction of cooperation when increasing *r* corresponds to a critical phase transition that follows directed percolation in both cases [26]. This can be interpreted in the sense that under harsh conditions, large *r*, the importance for cooperators to form compact clusters and reduce exploitation by defectors outweighs competition between members of the two sub-lattices.

### A. Symmetry breaking

At lower *r*, spontaneous symmetry breaking in the levels of cooperation on each sub-lattice marks another critical phase transition that belongs to the universality class for spontaneous magnetization in the Ising model. As a result, players in one sub-lattice exploit those in the other, on average. However, in any asymmetric dynamical equilibrium, exploitation is ephemeral and limited both in time and space. Numerical simulations and analytical approximations suggest that spontaneous symmetry breaking occurs when the average cooperation across lattices becomes sufficiently large – approximately,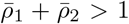. Similar spontaneous symmetry breaking was reported previously in two-layer games [54, 65]. More specifically, the competition between sub-populations is reminiscent of models for mutualistic interactions between species, where each species resides on their own lattice layer [32, 33]. In fact, the spatial inter-species donation game in [33]exhibits qualitatively the same dynamical behaviour as reported here, including the critical phase transitions, burst dynamics and symmetry breaking. However, the inter-species setup precludes analytical approaches to approximate and better understand the microscopic mechanisms that drive the macroscopic dynamics. Here we consider a different population structure that preserves the dynamical features but also admits approximations based on the four-site cluster method, which accurately predicts the qualitative dynamics.

### B. Effects of noise

In both models an optimal noise level in the updating procedure exists, but to very different ends: maximizing the range of *r* supporting cooperation for traditional overlapping neighbourhoods (see Fig. 6), while maximizing the asymmetry between sub-lattices for disjoint neighbourhoods (see Fig. 14). Optimal noise levels have previously been reported for the weak prisoner’s dilemma on lattices [11, 27, 66] as well as in spatial games with players using imitation rules that are characterized by different levels of noise [67]. Similar optimal noise levels are also discussed as coherence resonance [68–71] or stochastic resonance [72] and reflect the importance of *short-range* correlations. The impact of those correlations is suppressed with both low and high noise: either longer-range correlations become dominant, or short-range correlations get destroyed. In contrast, models that include self-interactions [27, 73] or possess connectivity structures that support the expansion of cooperation through overlapping small cliques (e.g., Moore neighbourhood on square lattice or triangular lattice with nearest neighbour interactions) on the two-dimensional systems do not exhibit such optima [11, 66, 67].

The asymmetry of cooperation between sub-lattices, 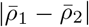, increases for decreasing cost-to-benefit ratio *r* and reaches a maximum. For still smaller *r*, bursts of defection and cyclical invasions of spatial strategy associations are the drivers of large fluctuations, which effectively decrease the asymmetry. Finally, for very small *r* the population occasionally ends in homogeneous defection but usually reaches absorbing states with one sub-lattice cooperating and the other defecting, and thus reflects perfect asymmetry.

### C. Spite

The persistence of spiteful behaviour, *r <* 0, is only observed in the traditional setup. The dynamics of the defector distribution mimics a branching-coalescing-annihilating random walk [42, 43], with exceedingly long coalescence times and vanishing probabilities to eliminate the last defector. The extinction of defectors exhibits a critical phase transitions that again falls into the universality class of directed percolation, see Fig. 4. The persistence of spiteful behaviour turns out to be an anomaly of the von Neumann neighbourhood because the clustering coefficient is zero and is not observed for other neighbourhoods, such as the Moore neighbourhood (*k* = 8), or on other lattices [11]. For disjoint sub-lattices, spite cannot be sustained and is replaced by homogeneous cooperation – as expected for interactions where cooperation is the dominant strategy.

### D. Asymmetric cooperation

In spatial models the four locally stable configurations of *CC, CD, DC*, and *DD* domains, can be viewed as strategy associations that compete against each other via complex invasion processes [11]. The larger number of locally stable stationary states, and transitions between them, gives rise to richer dynamics and, in particular, the emergence of spontaneous symmetry breaking in the frequency of cooperation in sub-populations that each reside on their own sub-lattice.

Competition between sub-populations has the surprising effect that under more benign conditions for cooperation, smaller *r*, asymmetric outcomes with members of one sub-lattice exploiting those of the other are favoured. The asymmetry tends to increase further for decreasing *r* and eventually results in fully asymmetric absorbing configurations.

## ACKNOWLEDGMENTS

C.H. acknowledges financial support from Natural Sciences and Engineering Research Council of Canada (NSERC) RGPIN-2021-02608.

## References

[1] J. Maynard Smith, Evolution and the Theory of Games (Cambridge University Press, Cambridge, UK, 1982).

[2] J. Hofbauer and K. Sigmund, The Theory of Evolution and Dynamical Systems (Cambridge University Press, Cambridge, UK, 1988).

[3] J. W. Weibull, Evolutionary Game Theory (MIT Press, Cambridge, MA, 1995).

[4] M. A. Nowak, Evolutionary Dynamics (Harvard University Press, Cambridge, MA, 2006).

[5] E. Frey, Physica A 389, 4265 (2010).

[6] J. Tanimoto, Fundamentals of evolutionary game theory and its applications (Springer, Tokyo, 2015).

[7] M. Perc, J. J. Jordan, D. G. Rand, Z. Wang, S. Boccaletti, and A. Szolnoki, Phys. Rep. 687, 1 (2017).

[8] V. Capraro and M. Perc, J. R. Soc. Interface 18, 20200880 (2021).

[9] A. Traulsen and N. E. Glynatsi, Phil. Trans. R. Soc. B 378, 20210508 (2023).

[10] M. A. Nowak and R. M. May, Nature 359, 826 (1992).

[11] G. Szabò and G. Fáth, Phys. Rep. 446, 97 (2007).

[12] C. P. Roca, J. A. Cuesta, and A. Sánchez, Phys. Life Rev. 6, 208 (2009).

[13] K. Sigmund and M. A. Nowak, Current Biology 9, R503 (1999).

[14] K. Sigmund, Theor. Popul. Biol 68, 7 (2005).

[15] C. Hauert, F. Michor, M. A. Nowak, and M. Doebeli, J. Theor. Biol 239, 195 (2006).

[16] R. M. Dawes, Ann. Rev. Psychol. 31, 169 (1980).

[17] K. Sigmund, The Calculus of Selfishness (Princeton University Press, Princeton, NJ, 2010).

[18] W. D. Hamilton, Nature 228, 1218 (1970).

[19] W. L. Vickery, J. S. Brown, and G. J. FitzGerald, Oikos 102, 413 (2003).

[20] T. M. Liggett, Stochastic interacting systems: Contact, voter and exclusion processes (Springer, New York, 1991).

[21] R. Axelrod, The Evolution of Cooperation (Basic Books, New York, 1984).

[22] R. Sugden, The economics of rights, cooperation and welfare (Basil Blackwell, Oxford, 1986).

[23] C. Hauert and M. Doebeli, Nature 428, 643 (2004).

[24] M. Doebeli and C. Hauert, Ecol. Lett. 8, 748 (2005).

[25] H. Ohtsuki, C. Hauert, E. Lieberman, and M. A. Nowak, Nature 441, 502 (2006).

[26] G. Szabò and C. Hauert, Phys. Rev. Lett. 89, 118101 (2002).

[27] G. Szabò and C. Tãke, Phys. Rev. E 58, 69 (1998).

[28] H. Ohtsuki, J. M. Pacheco, and M. A. Nowak, J. Theor. Biol. 246, 681 (2007).

[29] H. Ohtsuki, M. A. Nowak, and J. M. Pacheco, Phys. Rev. Lett. 98, 108106 (2007).

[30] M. Ifti, T. Killingback, and M. Doebeli, J. Theor. Biol. 231, 97 (2004).

[31] C. Hauert, EvoLudo: Evolutionary dynamics simulation toolkit (2025), URL https://www.evoludo.org.

[32] M. Doebeli and N. Knowlton, Proc. Natl. Acad. Sci. USA 95, 8676 (1998).

[33] C. Hauert and G. Szabò, PNAS Nexus 3, pgae326 (2024).

[34] H. E. Stanley, Introduction to Phase Transitions and Critical Phenomena (Clarendon Press, Oxford, 1971).

[35] H. A. Gutowitz, J. D. Victor, and B. W. Knight, Physica D 28, 18 (1987).

[36] R. Dickman, Phys. Rev. A 38, 2588 (1988).

[37] G. Szabò, A. Szolnoki, and R. Izsák, J. Phys. A: Math. Gen. 37, 2599 (2004).

[38] H. K. Janssen, Z. Phys. B 42, 151 (1981).

[39] P. Grassberger, Z. Phys. B 47, 365 (1982).

[40] J. Marro and R. Dickman, Nonequilibrium Phase Transitions in Lattice Models (Cambridge University Press, Cambridge, UK, 1999).

[41] H. Hinrichsen, Adv. Phys. 49, 815 (2000).

[42] T. M. Liggett, Interacting Particle Systems (Springer, New York, 1985).

[43] J. L. Cardy and U. C. Taüber, J. Stat. Phys. 90, 1 (1998).

[44] C. Hauert, Proc. R. Soc. Lond. B 268, 761 (2001).

[45] P. Clifford and A. Sudbury, Biometrika 60, 581 (1973).

[46] E. Ben-Naim, L. Frachebourg, and P. L. Krapivsky, Phys. Rev. E 53, 3078 (1996).

[47] G. F. Newell and E. W. Montroll, Rev. Mod. Phys. 25, 353 (1953).

[48] S. G. Brush, Rev. Mod. Phys. 39, 883 (1967).

[49] C. Domb, in Phase Transitions and Critical Phenomena, Vol. 3, edited by C. Domb and M. S. Green (Academic Press, London, 1974), pp. 357–484.

[50] P. C. Hohenberg and B. I. Halperin, Rev. Mod. Phys. 49, 435 (1977).

[51] Y. Lin and F. Wang, Phys. Rev. E 93, 022113 (2016).

[52] Q. Jin, L. Wang, C.-Y. Xia, and Z. Wang, Sci. Rep. 4, 4095 (2014).

[53] H. Lugo, J. C. González-Avella, and M. S. Miguel, Chaos 30, 083125 (2020).

[54] T. Takesue, Appl. Math. Comput. 388, 125543 (2021).

[55] B. Gao, J. Hong, H. Guo, S. Dong, and Z.-Z. Lan, Physica A 609, 128320 (2023).

[56] J. D. Gunton and M. S. Miguel, in Phase Transitions and Critical Phenomena, Vol. 8, edited by C. Domb and J. L. Lebowitz (Academic Press, New York, 1983).

[57] A. J. Bray, Adv. Phys. 43, 357 (1994).

[58] R. Hanel, S. A. Kauffman, and S. Thurner, Phys. Rev. E 76, 036110 (2007).

[59] Z. Yang and Z. Li, Nonlin. Dyn. 108, 4599 (2022).

[60] K. Binder and H. Müller-Krumbhaar, Phys. Rev. B 9, 2328 (1974).

[61] D. W. Oxtoby, J. Phys.: Condens. Matter 4, 7627 (1992).

[62] K. Hòdsági and G. Szabò, Physica A 525, 1379 (2019).

[63] Z.-X. Wu and Y.-H. Wang, Phys. Rev. E 75, 041114 (2007).

[64] J. Zhang, C. Zhang, T. Chu, and F. J. Weissing, PLoS ONE 9, e90288 (2014).

[65] H. Ezoe, J. Theor. Biol. 259, 744 (2009).

[66] G. Szabò, J. Vukov, and A. Szolnoki, Phys. Rev. E 72, 047107 (2005).

[67] G. Szabò, A. Szolnoki, and J. Vukov, EPL 87, 18007 (2009).

[68] A. S. Pikovsky and J. Kurths, Phys. Rev. Lett. 78, 775 (1997).

[69] O. Carrillo, M. A. Santos, J. Garcia-Ojalvo, and J. M. Sancho, EPL 65, 452 (2004).

[70] M. Perc and M. Marhl, New J. Phys. 8, 142 (2006).

[71] J. Tanimoto, Physica A 462, 714 (2016).

[72] L. Gammaitoni, P. Hänggi, P. Jung, and F. Marchasoni, Rev. Mod. Phys. 70, 223 (1998).

[73] M. A. Nowak and R. M. May, Int. J. Bifurcat. Chaos 3, 35 (1993).

